# Multiple instance learning to predict immune checkpoint blockade efficacy using neoantigen candidates

**DOI:** 10.1101/2022.05.06.490587

**Authors:** Franziska Lang, Patrick Sorn, Barbara Schrörs, David Weber, Stefan Kramer, Ugur Sahin, Martin Löwer

**Affiliations:** TRON Translational Oncology gGmbH, Mainz, Germany; Institute of Computer Science, Johannes Gutenberg University, Mainz, Germany; BioNTech SE, Mainz, Germany; University Medical Center of the Johannes Gutenberg University, Mainz, Germany

**Author notes:** Corresponding author: Martin Löwer.

## Abstract

A successful response to immune checkpoint blockade treatment (ICB) depends on the functional re-invigoration of neoantigen-specific T cells and their anti-tumoral activity. Previous studies showed that the patient’s neoantigen candidate load is an imperfect predictor of the response to ICB. Further studies provided evidence that the overall response to ICB is also affected by the qualitative properties of a few or even single candidates, limiting the predictive power based on candidate quantity alone.

To our knowledge, this is the first study to predict the response to ICB therapy based on qualitative neoantigen candidate profiles in the context of the mutation type, using a multiple instance learning approach. Multiple instance learning is a special branch of machine learning which classifies labelled bags that are formed by a set of unlabeled instances. The multiple instance learning approach performed systematically better than random guessing and was independent of the neoantigen candidate load. Qualitative modeling performed better in comparison to the quantitative approach, in particular for modelling low-abundant fusion genes. Our findings suggest that multiple instance learning is an appropriate method to predict immunotherapy efficacy based on qualitative neoantigen candidate profiles without relying on direct T-cell response information and provide a foundation for future developments in the field.

## Introduction

Neoantigens are cancer-specific mutated gene products that are presented in the form of neoepitopes by the major histocompatibility complex (MHC) proteins and recognized by CD8^+^ or CD4^+^ T cells. Upon neoepitope recognition, these neoantigen-specific T cells can mediate tumor control in the presence of a favorable tumor microenvironment. Immune checkpoint blockade (ICB) drives tumor control via the functional re-invigoration of neoantigen-specific T cells^1–4^. We previously introduced a concept-based classification of neoantigens, classifying neoantigens that are recognized by such pre-existing re-invigorated T cells and that are predictive for the clinical benefit of ICB therapy as restrained neoantigens^5^.

Different types of mutation sources can generate neoantigens with diverse molecular characteristics. While neoantigens from single nucleotide variants (SNVs) usually cause a single amino acid substitution, INDELs (small insertions or deletions) or fusion genes can generate frameshift neoantigens with completely altered amino acid sequences. INDELs can generate immunogenic neoantigens^6,7^ and the INDEL burden correlates with the response to ICB in melanoma patients^8,9^. Furthermore, a head and neck cancer patient with clinical response to anti-PD-1 therapy harbored only one single immunogenic neoantigen from a fusion gene^10^. These observations suggest that neoantigens of all mutation types could act as restrained neoantigens. Previous studies investigating the characteristics of restrained neoantigens from SNVs have shown that clonality^11^, the difference in MHC-I binding affinity to the wild-type peptide (differential agretopicity index, DAI)^12^ and the ratio-based DAI in combination with the sequence similarity to epitopes from known pathogens^13^ correlated with survival upon ICB therapy. Therefore, the response to cancer immunotherapy is driven not only by neoantigen candidate quantity but also by their quality^14^.

Further neoantigen features and prioritization methods have been published, and we recently developed a toolbox called NeoFox^15^ to annotate neoantigen candidates with a variety of neoantigen features. An analysis of how these features characterize restrained neoantigens is still missing - in particular in the context of non-SNV mutation types.

Standardized, unbiased and systematic immunogenicity screenings of neoantigen candidates providing direct information about neoantigen-specific T-cell responses are still limited in their availability for such an *in silico* analysis. Therefore, we predicted neoantigen candidates from raw whole exome (WES) and RNA sequencing data from five ICB cohorts to examine if the clinical response can be predicted based on the characteristics of neoantigen profiles in the context of the mutation type.

While traditional supervised machine learning approaches classify labelled instances, multiple instance learning is a special branch which classifies labelled groups (so-called bags) that are formed by a set of instances with unknown labels^16^. According to the standard assumption of multiple instance learning, positive bags harbor at least one instance with a hidden positive label and negative bags harbor exclusively negative instances^16,17^. For our analysis, patients are referred to as bags and the clinical response to ICB is used as the bag label. Multiple instance learning has been used in the field of cancer immunology for distinguishing tumor from normal samples on their T-cell receptor (TCR) sequence profiles^18^ and for predicting T-cell infiltration on neoantigen candidate profiles^19^.

Here, we used multiple instance learning to predict the clinical response to ICB based on neoantigen candidate profiles of cancer patients in the context of the mutation type. We further identified features that are relevant to predict ICB efficacy and that may characterize restrained neoantigens.

## Methods

### Datasets and their collection

Independent datasets from five immune checkpoint blockade trials were collected^20–24^.

Whole exome sequencing (WES) from tumor and matched normal samples and RNA-seq data of the tumor sample were retrieved from the respective repositories.

Clinical outcome data was collected from the original publications, and only patients with both available WES and RNA-seq were considered in the downstream analysis. The response categories were transformed into a table of binary outcomes. Patients with complete (CR) or partial (PR) response were defined as responders and patients with stable (SD) or progressive (PD) disease as non-responders.

Samples were restricted to ICB-therapy naïve samples that were acquired pre-treatment in the Riaz cohort^22^. Only patients treated with atezolizumab as a single agent were considered in the analysis in the McDermott cohort^23^.

### Prediction of neoantigen candidates

Neoantigen candidates were detected using an in-house build standardized pipeline that was described previously^25,26^. The pipeline covers the alignment of DNA reads to the reference genome hg19 using bwa (v0.7.10)^27^ and the removal of duplicated reads with Picard (v1.110) (http://broadinstitute.github.io/picard). An in-house developed proprietary software was used to detect high-confidence single nucleotide variations. INDEL variations were detected with strelka2^28^. The detected somatic nonsynonymous mutations were translated into 27mer peptide sequences with the mutation at position 14. Frameshift INDELs were translated until the occurrence of the next stop codon. Unsolvable technical issues arose for two patients (Hugo: Pt8, McDermott: Pt164370145747331); these were excluded from further analysis that included SNV- or INDEL-derived neoantigen candidates.

Neoantigen candidates derived from fusion genes were predicted from RNA-seq data using EasyFuse^29^. For the downstream analyses, fusion gene-derived neoantigen candidates had to fulfil the following criteria: (i) an EasyFuse probability score > 0.5, (ii) to not be a common falsepositive fusion gene call (iii) best break point per fusion gene pair based on the prediction probability of the random forest classifier, (iv) breakpoints must be on the respective exon boundary, (v) frame is not “no_frame” and (vi) exclude neoantigen candidates with “neo_frame” in case “in_frame” neoantigen candidates were predicted for the same fusion gene.

### HLA-typing

MHC -I and –II genotypes were detected for each patient with HLA-HD (v1.2.0.1) using the normal WES-seq data^30^.

### Transcript expression analysis

Transcript expression analysis was performed by aligning RNA-seq reads to the hg19 reference genome with STAR (v2.4.2a)^31^, followed by quantification of transcripts in FPKM (fragments per kilobase of exon model per million reads mapped) with sailfish (vBeta-0.7.6)^32^.

### Annotation of neoantigen candidates

Neoantigen candidates from all mutation types were annotated with published neoantigen features and prioritization algorithms using the NeoFox (v0.5.3) toolbox^15^. The predicted neoantigen candidates, MHC-I and -II genotypes of the patient and the tumor type were provided as input.

The wild type counterpart for neoepitope candidates from INDELs or fusion genes was defined by the best hit of the same length in a BLAST (Basic Local Alignment Search Tool) search against the human proteome in NeoFox.

### Multiple Instance Learning

The response to ICB was predicted with multiple instance learning on the annotated neoantigen candidates using the MILES (Multiple-Instance Learning via Embedded Instance Selection) algorithm from the python library mil (https://github.com/rosasalberto/mil).

Twenty-nine neoantigen features from NeoFox were included in the analysis (Table 1). The direction of scaling was harmonized for all neoantigen features, i.e. the scaling was reversed for Best_rank_MHCI_score, Best_rank_MHCII_score, Dissimilarity_MHCI, MixMHC2pred_best_rank, MixMHCpred_best_rank, Selfsimilarity_MHCI, Selfsimilarity_MHCII, PRIME_best_rank, PHBR_I, PHBR_II. Missing values were filled with the minimal value of a neoantigen feature across all predicted neoantigen candidate, assuming that a missing value reflects biological irrelevance of the neoantigen candidate of interest. Data was scaled using the “StandarizerBagsList” function of the mil library.

**Table 1.**

| Feature | Description | Reference |
| --- | --- | --- |
| rnaExpression | RNA expression | - |
| rnaVariantAlleleFrequency | Variant allele fraction | - |
| Best_rank_MHCI_score | Best predicted MHC-I binding rank per neoantigen candidate | 46 |
| Best_rank_MHCII_score | Best predicted MHC-II binding rank per neoantigen candidate | 46 |
| MixMHCpred_best_rank | Best predicted MixMHCpred rank score per neoantigen | 47 |
| MixMHC2pred_best_rank | Best predicted MixMHC2pred rank score per neoantigen | 48 |
| Amplitude_MHCI_affinity | Ratio of the MHC-I affinity score between the best predicted MHC-I neoepitope and its corresponding WT peptide | 13,41 |
| Amplitude_MHCII_rank | Ratio of the MHC-II rank score between the best predicted MHC-II neoepitope and its corresponding WT peptide | Adapted from<sup>13,41</sup> |
| DAI_MHCI_affinity | Difference in the MHC-I affinity score between the best predicted MHC-I neoepitope and its corresponding WT peptide | 40 |
| PHBR_I | The harmonic mean of best predicted MHC-I binding rank across the MHC-I genotype | 49 |
| PHBR_II | The harmonic mean of best predicted MHC-II binding rank across the MHC-II genotype | 50 |
| Generator_rate_MHCI | Number of predicted MHC-I neoepitopes per neoantigen | 51 |
| Generator_rate_MHCII | Number of predicted MHC-II neoepitopes per neoantigen | 51 |
| Selfsimilarity_MHCI | Similarity to the self-proteome of the best predicted MHC-I neoepitope per neoantigen candidate | 52 |
| Selfsimilarity_MHCII | Similarity to the self-proteome of the best predicted MHC-II neoepitope per neoantigen candidate | Adapted from<sup>52</sup> |
| Dissimilarity_MHCI | Similarity to the self-proteome of the best predicted MHC-I neoepitope per neoantigen candidate | 53 |
| Pathogensimilarity_MHCI_9mer | Similarity of the best predicted MHC-I neoepitope per neoantigen candidate to known pathogens | 13,41 |
| Pathogensimilarity_MHCII | Similarity of the best predicted MHC-II neoepitope per neoantigen candidate to known pathogens | Adapted from<sup>13,41</sup> |
| Hex_alignment_score_MHCI | Similarity of the best predicted MHC-I neoepitope per neoantigen candidate to known pathogens | 39 |
| Hex_alignment_score_MHCII | Similarity of the best predicted MHC-II neoepitope per neoantigen candidate to known pathogens | Adapted from<sup>39</sup> |
| IEDB_Immunogenicity_MHCI | IEDB Immunogenicity score for the best predicted MHC-I neoepitope | 54 |
| IEDB_Immunogenicity_MHCII | IEDB Immunogenicity score for the best predicted MHC-II neoepitope | Adapted from<sup>54</sup> |
| vaxrank_binding_score | Cumulative MHC I binding score per neoantigen candidate | 38 |
| vaxrank_total_score | Combination of vaxrank binding score with variant allele expression | 38 |
| Priority_score | Combinatorial score of MHC-I binding rank and variant allele expression | 55 |
| Recognition_Potential_MHCI_9mer | Combinatorial score of amplitude MHC-I and pathogensimilarity | 41 |
| **Neoag_immunogenicity** | Machine Learning model | 13,56 |
| **Tcell_predictor_score** | Machine Learning model | 57 |
| **PRIME_best_rank** | Machine Learning model | 58 |

Models were trained with a plain 10-fold cross validation on the full dataset to allow a robust estimation of the performance of the learning method across the repeated splits^33^.

The MILES algorithm comes with the two hyperparameters sigma2 and c ^34^. To set the hyperparameters (c=[0.1,…,1], sigma2=[50,…, 10000000]), an internal 10-fold cross validation was used. Thus, a 10-fold external cross validation was used for model validation and within each “fold” another internal 10-fold cross-validation for hyperparameter estimation, amounting to in total 100 runs in a nested cross-validation.

This approach was used to estimate the performance of the learning method in predicting the response to ICB on neoantigen candidates restricted to SNVs, INDELs or fusion genes or on a combined dataset covering neoantigen candidates from SNVs, INDELs and fusion genes. The performance of the learning method was estimated by the area under the receiver operating characteristic curve (AUROC).

To estimate the importance of each neoantigen feature, the feature of interest was permutated and models were re-trained on that dataset using the best hyper-parameter setting of the original approach.

## Results

### Neoantigen candidate profiles are heterogeneous in cancer patients

To investigate the characteristics of neoantigen candidates in the context of their mutation type, we identified neoantigen candidates from SNVs, INDELs and fusion genes in raw WES and RNA-seq data from five melanoma or renal cell carcinoma patient cohorts treated with α-PD-1^21,22,24^, α-CTLA-4^20^ or α-PD-L1^23,24^ cancer immunotherapy. We then annotated these neoantigen profiles with their characteristics (Figure 1).

**Figure 1:**
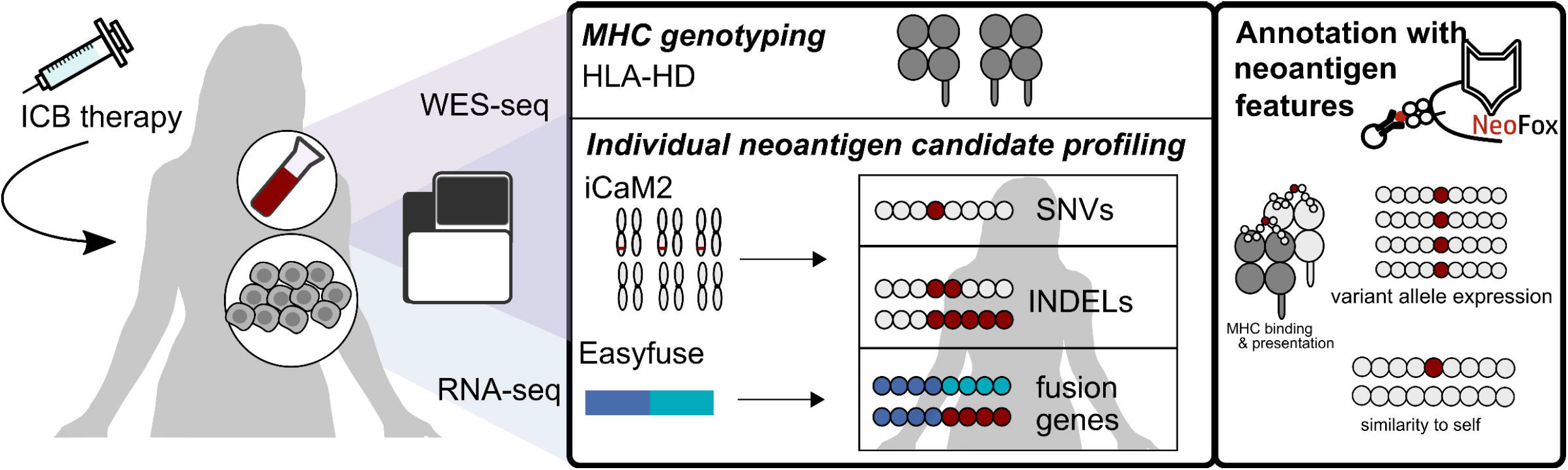
Identification of neoantigen candidates in immune checkpoint blockade (ICB) cohorts. Publicly available WES and RNA-seq from patient cohorts who had been treated with ICB therapy were collected. Neoantigen candidates from SNVs and INDELs were identified with an in-house property pipeline^25^ (“iCaM2”) and candidates from fusion genes were identified with EasyFuse^29^. Neoantigen candidates were annotated with features and subjected to prioritization algorithms using NeoFox^15^.

The number of predicted neoantigen candidates varied between mutation types and patient cohorts. While the average load of neoantigen candidates derived from SNVs was 370 in the three melanoma cohorts (“Hugo”, “Riaz”, “Van Allen”), the average SNV-derived load was 77 in the two renal cell carcinoma cohorts (“Miao”, “McDermott”) (Figure 2A). Neoantigen candidates from INDELs or fusion genes were in general rarer than SNV-derived neoantigen candidates. The proportion of neoantigen candidates from INDELs or fusion genes was higher in patients of the renal cell carcinoma cohorts (“RCC”) in comparison to the melanoma cohorts (“MEL”) (Figure 2B).

**Figure 2:**
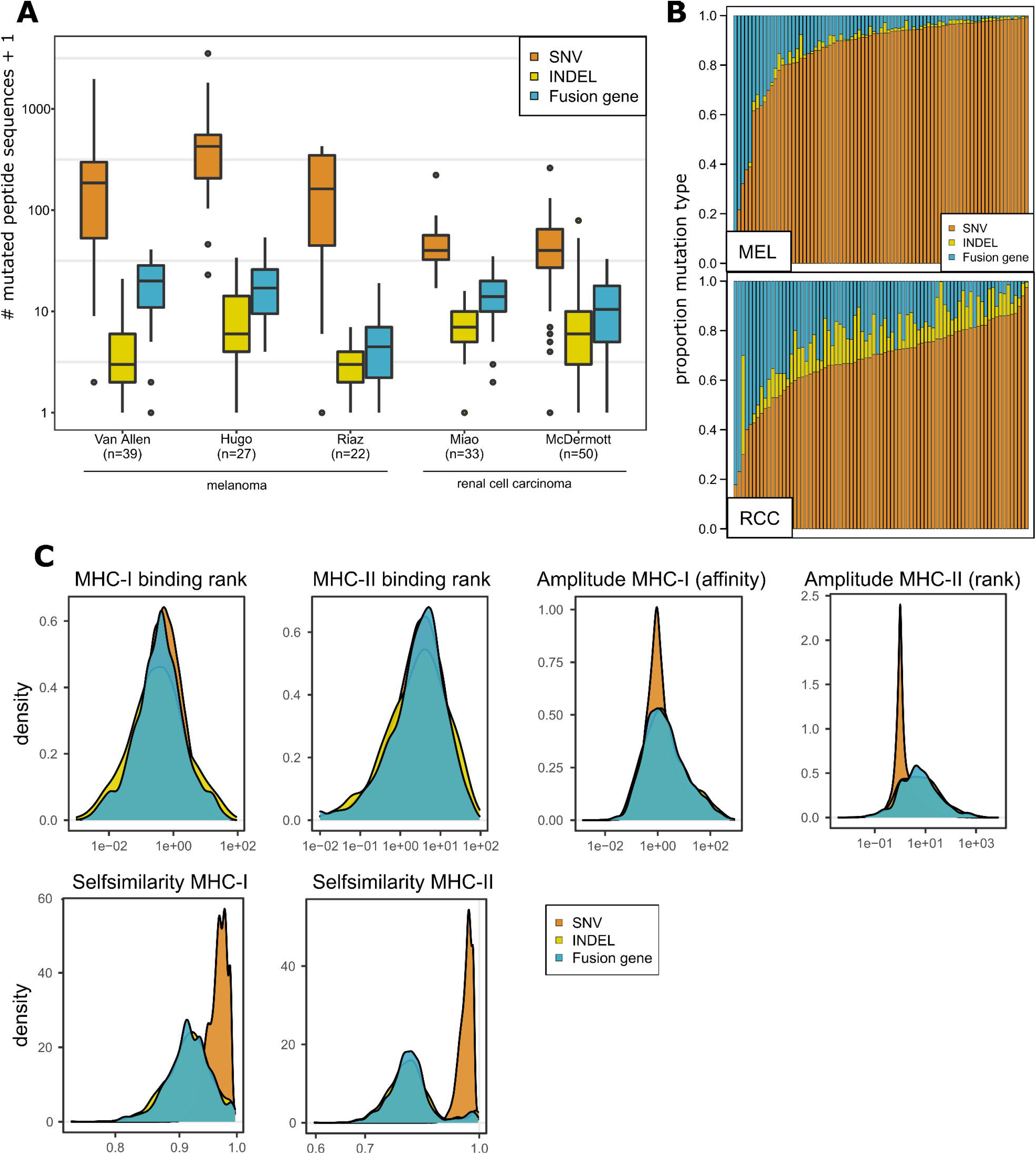
Neoantigen candidate profiles in cohorts treated with immune checkpoint blockade (ICB). (**A**) The number of neoantigen candidates derived from SNVs, INDELs and fusion genes in five ICB cohorts. Data are shown as median +/- interquartile range. (**B**) The relative composition of neoantigen candidate profiles with respect to mutation type was compared between the melanoma cohorts (“MEL”) and the renal cell carcinoma cohorts (“RCC”). (**C**) Density plots comparing the distribution of MHC-I binding rank, MHC-II binding rank, amplitude MHC-I, amplitude MHC-II, self-similarity MHC-I and self-similarity MHC-II between neoantigen candidates of different origin in the combined dataset of all ICB cohorts.

Next, we combined neoantigen candidates from the five ICB cohorts and compared the distribution of neoantigen characteristics between SNV, INDEL and fusion gene derived candidates (Figure 2C). The best-predicted MHC-I and MHC-II epitopes of SNV, INDEL and fusion gene derived neoantigen candidates shared comparable MHC binding ability. INDELs and fusion genes were associated with higher amplitude MHC-II (rank) values in comparison to SNVs. This suggested that the non-SNV mutation types are more likely to generate predicted MHC-II epitopes with improved MHC-II binding ranks compared to their wild type counterpart. As expected, the best-predicted MHC-I and MHC-II epitopes of INDEL- and fusion gene-derived neoantigen candidates were less similar to their wild type counterpart in comparison to SNV-derived candidates.

### The neoantigen candidate load is an imperfect predictor of the response to ICB

For each mutation type, we systematically evaluated whether the predicted neoantigen candidate load correlated with the response to ICB. The neoantigen candidate load was defined with respect to different MHC-I and MHC-II binding affinity thresholds, while considering either all or only expressed mutated peptide sequences (Figure 3A-D, Supplemental Figure 1A-J). The SNV-derived neoantigen candidate load was a general predictor of ICB efficacy when combining the melanoma cohorts (“MEL”) or all analyzed ICB cohorts (Figure 3D, Supplemental Figure 1A-C, J). The load of INDEL-derived neoantigen candidates with MHC-I or MHC-II binding ability < 50nM significantly predicted the response to ICB in the dataset covering all ICB cohorts (Figure 3D, Supplemental Figure 1D-F, J). In the melanoma cohort, also the load of all predicted neoantigen candidates from INDEL mutations was predictive with and without MHC binding pre-filtering (Figure 3D, Supplemental Figure 1J). Interestingly, the correlation of INDEL derived neoantigen candidates with good MHC-I or MHC-II binding properties was also apparent in one individual melanoma (“Riaz”) cohort (Figure 3A-D, Supplemental Figure 1J). The fusion gene derived neoantigen candidate load generally did not correlate with ICB efficacy and the load of neoantigen candidates from all three mutation types could be associated with response to ICB therapy but on lower significance levels compared to the SNV model (Figure 3D, Supplemental Figure 1J).

**Figure 3:**
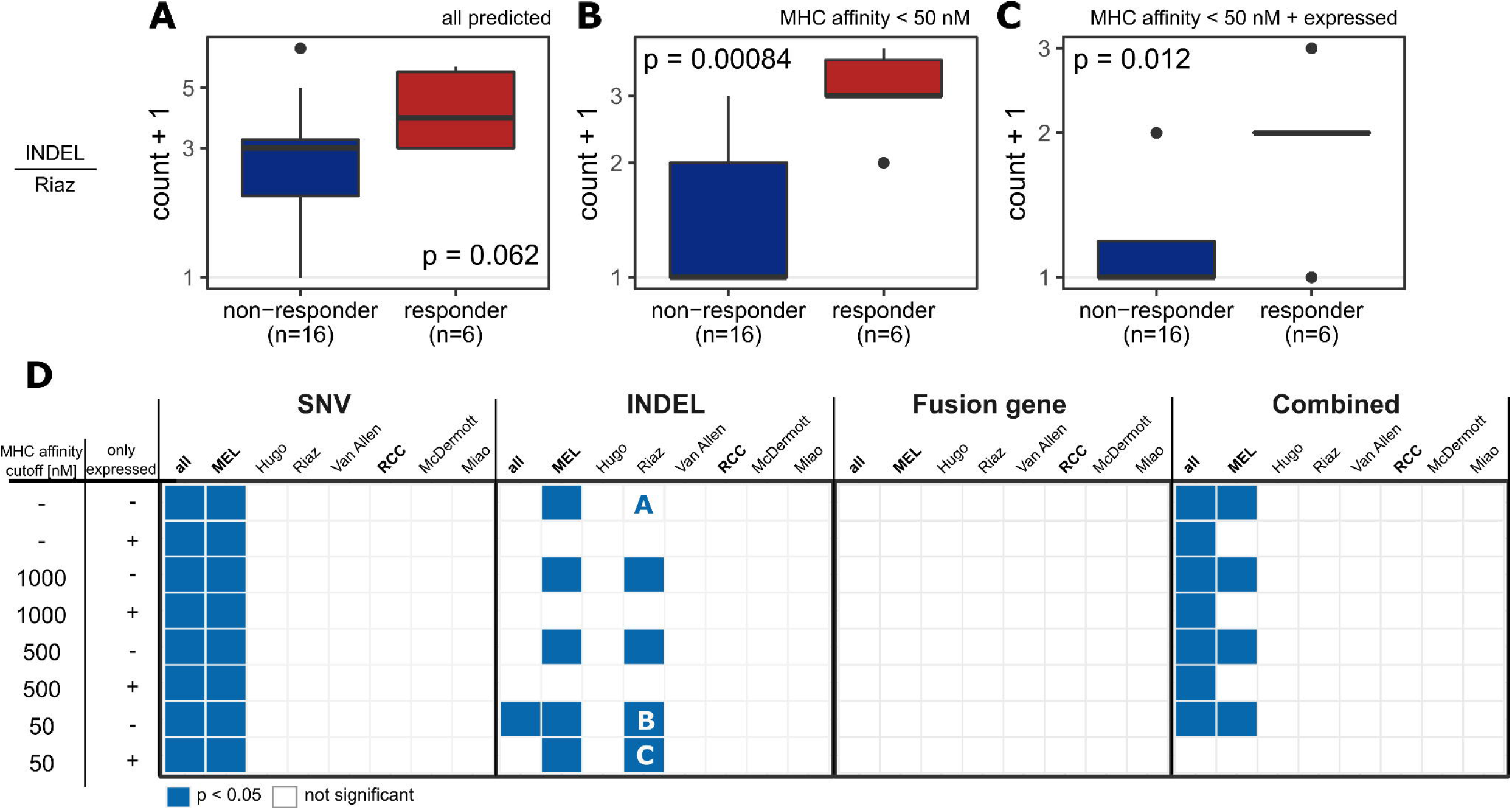
Neoantigen candidate load is an imperfect predictor of the response to ICB. **(A-C)** The INDEL-derived neoantigen candidate burden was compared between responder and non-responder the Riaz cohort based on (**A**) all predicted neoantigen candidates, (**B**) candidates with MHC-I or MHC-II binding affinity < 50 nM and (**C**) expressed candidates with MHC-I or MHC-II binding affinity < 50 nM. (**D**) The neoantigen candidate load was compared between responder and non-responder with respect to the mutation type, MHC binding ability and RNA expression. The plot represents the resulting the p-value from each comparison. Comparisons that resulted in a p-value < 0.05 are shown in blue while non-significant comparisons are shown in white. The y-axis represents a definition of the neoantigen candidate load. Neoantigen candidates with MHC-I or MHC-II binding affinity lower than the respective threshold (“MHC affinity cutoff”) were used. The column “only expressed” indicates if all neoantigen candidates or only neoantigen candidates confirmed in the RNA-seq (“+”) were used. All comparisons were performed in each individual ICB cohort and in a combined dataset of all cohorts (“all”), the melanoma cohorts (“MEL”) or the renal cell carcinoma cohorts (“RCC”). The letters A-C refer to respective subpanel above. Statistical testing was performed with Wilcoxon signed ranked test.

These observations support previous findings by other studies^35–37^ that the SNV or SNV-derived neoantigen candidate load alone is an imperfect predictor of the response to ICB.

### Multiple-Instance Learning via Embedded Instance Selection to predict the response to ICB on neoantigen candidate profiles

The imperfect correlation between neoantigen candidate load and the response to ICB motivated us to examine whether the response to ICB can be predicted by neoantigen candidate quality.

Therefore, we used multiple instance learning to predict the ICB efficacy based on a set of unlabeled neoantigen candidate profiles. Patients are referred to as bags with the response to ICB as their label (Figure 4A). Each bag is a collection of unlabeled instances, i.e. neoantigen candidates with unknown anti-tumoral activity. The multiple instance learning standard assumption meets the biological assumptions that responders (“positive bags”) must harbor at least one true neoantigen (“positive instance”) while non-responders (“negative bags”) harbor only neoantigen candidates that cannot trigger anti-tumoral activity (“negative instances”)^17^. The MILES (Multiple-Instance Learning via Embedded Instance Selection)^34^ algorithm was chosen as the algorithm of choice in this study as it performed well in a previous benchmarking study related to cancer detection based on TCR sequences^18^.

**Figure 4:**
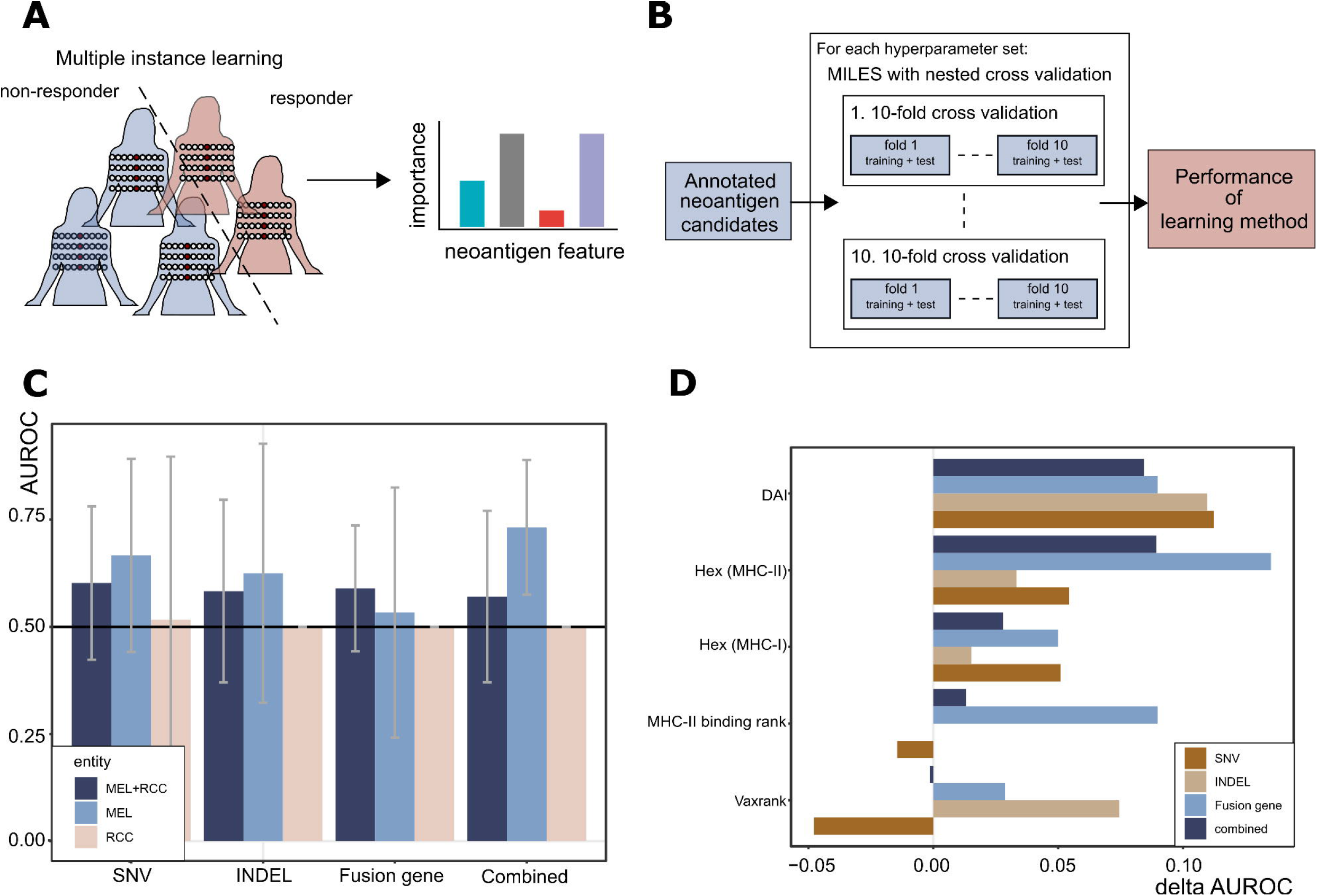
Multiple-Instance Learning via Embedded Instance Selection to predict the response to immune checkpoint blockade (ICB) using neoantigen candidates. (**A**) Multiple instance learning was used to distinguish responders from non-responders given their neoantigen candidate profiles as represented by the annotated neoantigen features. Upon identification of the best hyperparameter setting, the relevance of the individual neoantigen features is analyzed. (**B**) Models are trained and evaluated in a nested cross validation (CV) loop for multiple hyperparameter set on the full dataset. The hyperparameter set with the best median area under the receiver operating characteristic curve (AUROC) across the nested CV approach is selected to represent the performance of the learning method. (**C**) The median AUROC of MILES on a dataset of SNVs, Fusion genes or INDELS only and a dataset combining all mutation types (“combined”) for all cohorts (“MEL+RCC”), the melanoma cohorts (“MEL”) and the renal cell carcinoma cohorts (“RCC”). Data are shown as median +/- interquartile range over a nested CV. (**D**) To estimate the importance of each neoantigen features, the nested CV approach was repeated for the best hyperparameter setting on a dataset of all ICB cohorts in which the neoantigen feature of interest was permutated. Feature importance was approximated by the delta AUROC of learning method on the original data and the learning method on the data with permutated feature. Features with delta AUROC ≥0.05 in at least one approach are shown.

Neoantigen candidate profiles were encoded by 29 neoantigen features and prioritization methods such as MHC binding, transcript expression or the self-similarity (Table 1). We predicted the response to ICB on neoantigen candidates from SNVs, INDELs or fusion genes separately or by a combined set and analyzed the feature importance for each learning approach (Figure 4A).

For a robust performance estimation, the MILES algorithm was trained and evaluated on neoantigen candidates with a nested cross validation approach across multiple hyperparameter sets^33^ (Figure 4B). The median AUROC across the nested cross validation was used to evaluate the performance of the learning method. The set of hyperparameters with the best performance differed for the learning approaches trained on SNV, INDEL, fusion genes or on a combined set of neoantigen candidates (Supplemental Figure 2A).

Training and evaluating the MILES approach on SNV-derived neoantigen candidates from all ICB cohorts (“MEL+RCC”) achieved a median AUROC of 0.60 (Figure 4C, Supplemental Table 1). Similarly, the performance of MILES was better than random for fusion genes (median AUROC=0.59), INDELs (median AUROC = 0.58) and combined (median AUROC = 0.57). We observed an even better performance when evaluating MILES on a dataset restricted to the three melanoma cohorts (“MEL”) for the INDEL-specific (median AUROC = 0.63), SNV-specific (median AUROC = 0.67) and combined (median AUROC = 0.73) approach (Figure 4C, Supplemental Table 1). To compare the qualitative to the quantitative approach, we performed a ROC-curve analysis in a nested CV on the neoantigen candidate load as a predictor of ICB efficacy. This analysis suggests that the qualitative performed superior to the quantitative approach in particular for the combined (“MEL”, Supplemental Figure 2C) and fusion genespecific (“MEL+RCC”, Supplemental Figure 2D) data set (Supplemental Table 1).

As a control, we trained the MILES algorithm on datasets with randomized neoantigen candidates but original distribution of neoantigen candidate load (Supplemental Figure 2C-I). Here, MILES performed randomly (e.g. median AUROC = 0.53 for the SNV-specific approach in the “MEL+RCC” cohort:) indicating that prediction of the clinical response to ICB with MILES might indeed be driven by neoantigen quality and not by neoantigen quantity.

The MILES algorithm performed randomly on datasets restricted to data from renal cell carcinoma cohorts (“RCC”) (Figure 4C, Supplemental Table 1). The RCC cohorts had a higher fraction of patients with stable disease in comparison to the MEL cohorts (Supplemental Figure 3A). We hypothesized that stable disease leading e.g. to survival benefit may be mediated by restrained neoantigens and in the next analysis, excluded patients with stable disease. Of note, MILES achieved a median AUROC of 0.75 in the fusion gene-specific approach in RCC cohort, performing superior to the quantitative approach (Supplemental Figure 3B-D, Supplemental Table 1). MILES performed randomly on randomized neoantigen candidates in the RCC cohort (Supplemental Figure 3E-G).

Next, we examined which neoantigen features were important to predict ICB efficacy with MILES. Since the embedding step in the MILES algorithms involves a nonlinear transformation^34^, the feature importance could not be estimated internally within the algorithm. Therefore, we repeated the nested cross validation approach on datasets in which the neoantigen feature of interest was permutated. Then, we approximated feature importance by the delta AUROC of the original learning method and the approach without the feature of interest and considered features achieving a delta AUROC > 0.05 as relevant.

The results of the feature importance analysis were heterogeneous between the mutation types in the dataset covering all ICB cohorts (“MEL+RCC”) (Figure 4D). For instance, the MHC-II binding rank of best-predicted peptide was important only in the fusion gene-specific learning approach (delta AUROC = 0.09). Vice versa, vaxrank^38^ was predicted relevant in the case of INDELs (delta AUROC = 0.07) but not for SNVs or fusion genes. Interestingly, the similarity of the best-predicted MHC-II peptide to known pathogenic epitopes in terms of the HEX score^39^ achieved delta AUROCs higher than 0.05 for the learning approach on SNVs (delta AUROC = 0.05), fusion gene (delta AUROC = 0.14) and the combined dataset (delta AUROC = 0.09). The differential agretopicity index (DAI)^40^ was predicted as important feature in all approaches, suggesting its general relevance (Figure 4D, Supplemental Figure 4A, B). Furthermore, features such as the recognition potential^41^ (delta AUROC =0.08) and PRIME (delta AUROC = 0.08) were predicted to be relevant in the combined set of neoantigen candidates from all mutation types specifically in the context of RCC.

## Discussion

Predicting ICB therapy efficacy with neoantigens as drivers is still challenging due to lack of appropriate models. We tackled this challenge by comparing a quantitative and qualitative approach to predicting the response to ICB based on predicted neoantigen candidates from SNVs, INDELs and fusion genes.

In the quantitative approach, we found SNV-derived neoantigen candidate load as a predictor to the response to ICB in a combined set of all ICB cohorts studied but not in an individual ICB cohort. Our quantitative approach supported previous findings^8,9^ that the neoantigen candidate load from INDEL mutations correlates with the response to ICB in particular in the context of melanoma. Similar to previous observations^42^, we observed that the number of neoantigen candidates from fusion genes was not an indicator for the response to ICB. The limitations of the mutation or neoantigen candidate load as a predictor of the response to ICB has been extensively studied and discussed in particular in the context of SNVs^35–37^. It is conceivable that technical shortcomings in the tools used for mutation calling and in the definition of the neoantigen candidate load limit its predictive power in our and other studies. Nevertheless, the observation that patients with low neoantigen candidate load also harbor immunogenic neoantigens and can respond to ICB therapy^10^ suggests that, aside from technical shortcomings, disregarding qualitative traits limits and pursuing a purely quantitative approach would limit its predictive power.

Previous studies have used different approaches and prior assumptions to predict the response to ICB based on qualitative neoantigen profiles, e.g. based on the best-predicted neoantigen candidate^13,41^, the mean across all predicted candidates^12^ or by the Cauchy-Schwarz index^43^. However, to our knowledge, this is the first study that predicted the ICB therapy efficacy dependent on neoantigen profiles with multiple instance learning. This approach relies only on the prior assumption that a responder to ICB harbors at least one immunogenic neoantigen while a non-responder lacks immunogenic neoantigens. Evaluating MILES on neoantigen candidate data demonstrated its non-random performance and independence of the underlying neoantigen candidate load, as suggested by the evaluation of MILES on randomized data. The qualitative approach on the neoantigen candidates from fusion genes improved the prediction of clinical benefit, as compared with that based on the fusion gene load. In particular, predicting the ICB efficacy in RCC by fusion gene-derived neoantigen candidate profiles was superior to neoantigen candidate load when patients with stable disease were excluded from the analysis.

Previously, we defined neoantigens that are predictive of the clinical outcome of ICB therapy as restrained neoantigens^5^. We analyzed the relevance of neoantigen features for the learning method and confirmed a previous observation that the DAI of the best-predicted MHC-I neoepitope is a descriptor of neoantigen candidates from all mutation types that may contribute to ICB efficacy^12^. Furthermore, the similarity of the best-predicted MHC-II neoepitope to viral epitopes in terms of the HEX algorithm^39^ appeared to be a relevant neoantigen feature in our analysis. This observation indicates that at least a subset of the neoantigens in patients responding to ICB may be cross-recognized by heterologous T cells^39,44^. However, the external permutation based method to estimate feature importance comes with two main limitations: (i) feature importance results might change with permutation and (ii) some features might correlate and affect the importance measure of each other.

Here, we showed that multiple instance learning can be used to predict immunotherapy efficacy based on qualitative neoantigen candidate profiles covering multiple mutation types, and provide the basis for future investigation. Our results indicated that integrating the quantitative approach relying on the neoantigen candidate load and the qualitative multiple instance approach may improve the prediction of the response to ICB. Furthermore, a limited set of neoantigen features was integrated into the model approach in this study, mostly targeting the linear sequence of neoantigen candidates and rather focusing on the interaction with MHC molecules^15^. Integrating clonality information^11^, structural features^45^ and future features that specifically model the interaction between the MHC bound neoepitope candidate with the TCR repertoire may improve predictions. Additionally, when more data is available, systematic benchmarks may identify the best suitable multiple instance learning algorithm. However, one interesting characteristic of the MILES algorithm used in this study is its internal instance selection approach and its ability to be used for instance classification^34^. Therefore, multiple instance learning with instance selection could empower not just prediction of the clinical efficacy of ICB therapy but also the identification of immunogenic neoantigens in the future.

## Data availability

Data for the Hugo cohort^21^ is available in SRA under accession numbers SRP067938, SRP090294 (WES-seq) and SRP070710 (RNA-seq). Data for the Riaz cohort^22^ is available under accession numbers SRP095809 (WES-seq) and SRP094781 (RNA-seq). Data for the Van Allen cohort^20^ is available in dbGap under accession number phs000452.v2.p1. Data for the Miao cohort^24^ is available in dbGap under accession number phs001493.v1.p1. Data for the McDermott cohort^23^ is available in European Genome-Phenome Archive (EGA) under accession number EGAS00001002928. We acknowledge the authors and generators of these datasets and the grants that supported the studies.

Data including the predicted neoantigen candidates and code related to this study are available on Github (https://github.com/TRON-Bioinformatics/milneo_analysis).

## Acknowledgements

This work was supported by an ERC Advanced Grant to U.S. (ERC-AdG 789256). The authors would like to thank Karen Chu for proof-reading the manuscript and helpful comments.

## References

1. van Rooij, N. et al. Tumor exome analysis reveals neoantigen-specific T-cell reactivity in an ipilimumab-responsive melanoma. J. Clin. Oncol. 31, e439–e442 (2013).

2. Gubin, M. M. et al. Checkpoint blockade cancer immunotherapy targets tumour-specific mutant antigens. Nature 515, 577–581 (2014).

3. Snyder, A. et al. Genetic Basis for Clinical Response to CTLA-4 Blockade in Melanoma. N Engl J Med. 371, 2189–2199 (2014).

4. Alspach, E. et al. MHC-II neoantigens shape tumour immunity and response to immunotherapy. Nature 574, 696–701 (2019).

5. Lang, F., Schrörs, B., Löwer, M., Türeci, Ö. & Sahin, U. Identification of neoantigens for individualized therapeutic cancer vaccines. Nat Rev Drug Discov (2022).

6. Roudko, V. et al. Shared Immunogenic Poly-Epitope Frameshift Mutations in Microsatellite Unstable Tumors. Cell 183, 1634–1649.e17 (2020).

7. Cimen Bozkus, C. et al. Immune Checkpoint Blockade Enhances Shared Neoantigen-Induced T-cell Immunity Directed against Mutated Calreticulin in Myeloproliferative Neoplasms. Cancer Discov. 9, 1192–1207 (2019).

8. Litchfield, K. et al. Escape from nonsense-mediated decay associates with anti-tumor immunogenicity. Nat. Commun. 11, 3800 (2020).

9. Turajlic, S. et al. Insertion-and-deletion-derived tumour-specific neoantigens and the immunogenic phenotype. A pan-cancer analysis. Lancet Oncol. 18, 1009–1021 (2017).

10. Yang, W. et al. Immunogenic neoantigens derived from gene fusions stimulate T cell responses. Nat. Med. 25, 767–775 (2019).

11. McGranahan, N. et al. Clonal neoantigens elicit T cell immunoreactivity and sensitivity to immune checkpoint blockade. Science 351, 1463–1469 (2016).

12. Ghorani, E. et al. Differential binding affinity of mutated peptides for MHC class I is a predictor of survival in advanced lung cancer and melanoma. Ann. Oncol. 29, 271–279 (2017).

13. Łuksza, M. et al. A neoantigen fitness model predicts tumour response to checkpoint blockade immunotherapy. Nature 551, 517–520 (2017).

14. McGranahan, N. & Swanton, C. Neoantigen quality, not quantity. Sci. Transl. Med. 11, eaax7918 (2019).

15. Lang, F., Ferreiro, P. R., Löwer, M., Sahin, U. & Schrörs, B. NeoFox. Annotating neoantigen candidates with neoantigen features. Bioinformatics 37 (2021).

16. Dietterich, T. G., Lathrop, R. H. & Lozano-Pérez, T. Solving the multiple instance problem with axis-parallel rectangles. Artificial Intelligence 89, 31–71 (1997).

17. Foulds, J. & Frank, E. A review of multi-instance learning assumptions. The Knowledge Engineering Review 25, 1–25 (2010).

18. Xiong, D., Zhang, Z., Wang, T. & Wang, X. A comparative study of multiple instance learning methods for cancer detection using T-cell receptor sequences. Comput Struct Biotechnol J 19, 3255–3268 (2021).

19. Park, S. et al. Bayesian multiple instance regression for modeling immunogenic neoantigens. Stat Methods Med Res 29, 3032–3047 (2020).

20. van Allen, E. M. et al. Genomic correlates of response to CTLA-4 blockade in metastatic melanoma. Science 350, 207–211 (2015).

21. Hugo, W. et al. Genomic and Transcriptomic Features of Response to Anti-PD-1 Therapy in Metastatic Melanoma. Cell 165, 35–44 (2016).

22. Riaz, N. et al. Tumor and Microenvironment Evolution during Immunotherapy with Nivolumab. Cell 171, 934–949.e16 (2017).

23. McDermott, D. F. et al. Clinical activity and molecular correlates of response to atezolizumab alone or in combination with bevacizumab versus sunitinib in renal cell carcinoma. Nat. Med. 24, 749–757 (2018).

24. Miao, D. et al. Genomic correlates of response to immune checkpoint therapies in clear cell renal cell carcinoma. Science 359, 801–806 (2018).

25. Sahin, U. et al. Personalized RNA mutanome vaccines mobilize poly-specific therapeutic immunity against cancer. Nature 547, 222–226 (2017).

26. Hilf, N. et al. Actively personalized vaccination trial for newly diagnosed glioblastoma. Nature 565, 240–245 (2019).

27. Li, H. & Durbin, R. Fast and accurate short read alignment with Burrows-Wheeler transform. Bioinformatics 25, 1754–1760 (2009).

28. Kim, S. et al. Strelka2. Fast and accurate calling of germline and somatic variants. Nat. Methods 15, 591–594 (2018).

29. Weber, D. et al. Accurate detection of tumor-specific gene fusions reveals strongly immunogenic personal neo-antigens. Nat. Biotechnol. (2022).

30. Kawaguchi, S., Higasa, K., Shimizu, M., Yamada, R. & Matsuda, F. HLA-HD. An accurate HLA typing algorithm for next-generation sequencing data. Human Mutation 38, 788–797 (2017).

31. Dobin, A. et al. STAR. Ultrafast universal RNA-seq aligner. Bioinformatics 29, 15–21 (2012).

32. Patro, R., Mount, S. M. & Kingsford, C. Sailfish enables alignment-free isoform quantification from RNA-seq reads using lightweight algorithms. Nat. Biotechnol. 32, 462 EP - (2014).

33. Gütlein, M., Helma, C., Karwath, A. & Kramer, S. A Large-Scale Empirical Evaluation of Cross-Validation and External Test Set Validation in (Q)SAR. Mol Inform 32, 516–528 (2013).

34. Chen, Y., Bi, J. & Wang, J. Z. MILES. Multiple-instance learning via embedded instance selection. IEEE Trans Pattern Anal Mach Intell 28, 1931–1947 (2006).

35. Jardim, D. L., Goodman, A., Melo Gagliato, D. de & Kurzrock, R. The Challenges of Tumor Mutational Burden as an Immunotherapy Biomarker. Cancer Cell 39, 154–173 (2021).

36. Wood, M. A., Weeder, B. R., David, J. K., Nellore, A. & Thompson, R. F. Burden of tumor mutations, neoepitopes, and other variants are weak predictors of cancer immunotherapy response and overall survival. Genome Med. 12, 33 (2020).

37. McGrail, D. J. et al. High tumor mutation burden fails to predict immune checkpoint blockade response across all cancer types. Ann Oncol 32, 661–672 (2021).

38. Rubinsteyn, A. et al. Computational Pipeline for the PGV-001 Neoantigen Vaccine Trial. Front. Immunol. 8, 1807 (2018).

39. Chiaro, J. et al. Viral Molecular Mimicry Influences the Antitumor Immune Response in Murine and Human Melanoma. Cancer Immunol. Res. (2021).

40. Duan, F. et al. Genomic and bioinformatic profiling of mutational neoepitopes reveals new rules to predict anticancer immunogenicity. J. Exp. Med. 211, 2231–2248 (2014).

41. Balachandran, V. P. et al. Identification of unique neoantigen qualities in long-term survivors of pancreatic cancer. Nature 551, 512–516 (2017).

42. Wei, Z. et al. The Landscape of Tumor Fusion Neoantigens. A Pan-Cancer Analysis. iScience 21, 249–260 (2019).

43. Lu, T. et al. Tumor neoantigenicity assessment with CSiN score incorporates clonality and immunogenicity to predict immunotherapy outcomes. Sci. Immunol. 5, eaaz3199 (2020).

44. Leng, Q., Tarbe, M., Long, Q. & Wang, F. Pre-existing heterologous T-cell immunity and neoantigen immunogenicity. Clin. Transl. Immunology. 9, e01111 (2020).

45. Devlin, J. R. et al. Structural dissimilarity from self drives neoepitope escape from immune tolerance. Nat. Chem. Biol. 16, 1269–1276 (2020).

46. Reynisson, B., Alvarez, B., Paul, S., Peters, B. & Nielsen, M. NetMHCpan-4.1 and NetMHCIIpan-4.0. Improved predictions of MHC antigen presentation by concurrent motif deconvolution and integration of MS MHC eluted ligand data. Nucleic Acids Res. 48, W449–W454 (2020).

47. Bassani-Sternberg, M. et al. Deciphering HLA-I motifs across HLA peptidomes improves neoantigen predictions and identifies allostery regulating HLA specificity. PLoS Comput. Biol. 13, e1005725 (2017).

48. Racle, J. et al. Robust prediction of HLA class II epitopes by deep motif deconvolution of immunopeptidomes. Nat. Biotechnol. 37, 1283–1286 (2019).

49. Marty, R. et al. MHC-I Genotype Restricts the Oncogenic Mutational Landscape. Cell 171, 1272–1283.e15 (2017).

50. Marty Pyke, R. et al. Evolutionary Pressure against MHC Class II Binding Cancer Mutations. Cell 175, 416–428.e13 (2018).

51. Rech, A. J. et al. Tumor Immunity and Survival as a Function of Alternative Neopeptides in Human Cancer. Cancer Immunol. Res. 6, 276–287 (2018).

52. Bjerregaard, A.-M. et al. An Analysis of Natural T Cell Responses to Predicted Tumor Neoepitopes. Front. Immunol. 8, 1566 (2017).

53. Richman, L. P., Vonderheide, R. H. & Rech, A. J. Neoantigen Dissimilarity to the Self-Proteome Predicts Immunogenicity and Response to Immune Checkpoint Blockade. Cell Systems 9, 375–382.e4 (2019).

54. Calis, J. J. A. et al. Properties of MHC Class I Presented Peptides That Enhance Immunogenicity. PLoS Comput. Biol. 9, e1003266 (2013).

55. Bjerregaard, A.-M., Nielsen, M., Hadrup, S. R., Szallasi, Z. & Eklund, A. C. MuPeXI. Prediction of neo-epitopes from tumor sequencing data. Cancer Immunol. Immunother. 66, 1123–1130 (2017).

56. Smith, C. C. et al. Machine-Learning Prediction of Tumor Antigen Immunogenicity in the Selection of Therapeutic Epitopes. Cancer Immunol. Res. 7, 1591–1604 (2019).

57. Besser, H., Yunger, S., Merhavi-Shoham, E., Cohen, C. J. & Louzoun, Y. Level of neo-epitope predecessor and mutation type determine T cell activation of MHC binding peptides. J. Immunother. Cancer 7, 135 (2019).

58. Schmidt, J. et al. Prediction of neo-epitope immunogenicity reveals TCR recognition determinants and provides insight into immunoediting. Cell Reports Medicine 2, 100194 (2021).

