## Supplementary Information for "Multiple instance learning to predict immune checkpoint blockade efficacy using neoantigen candidates"

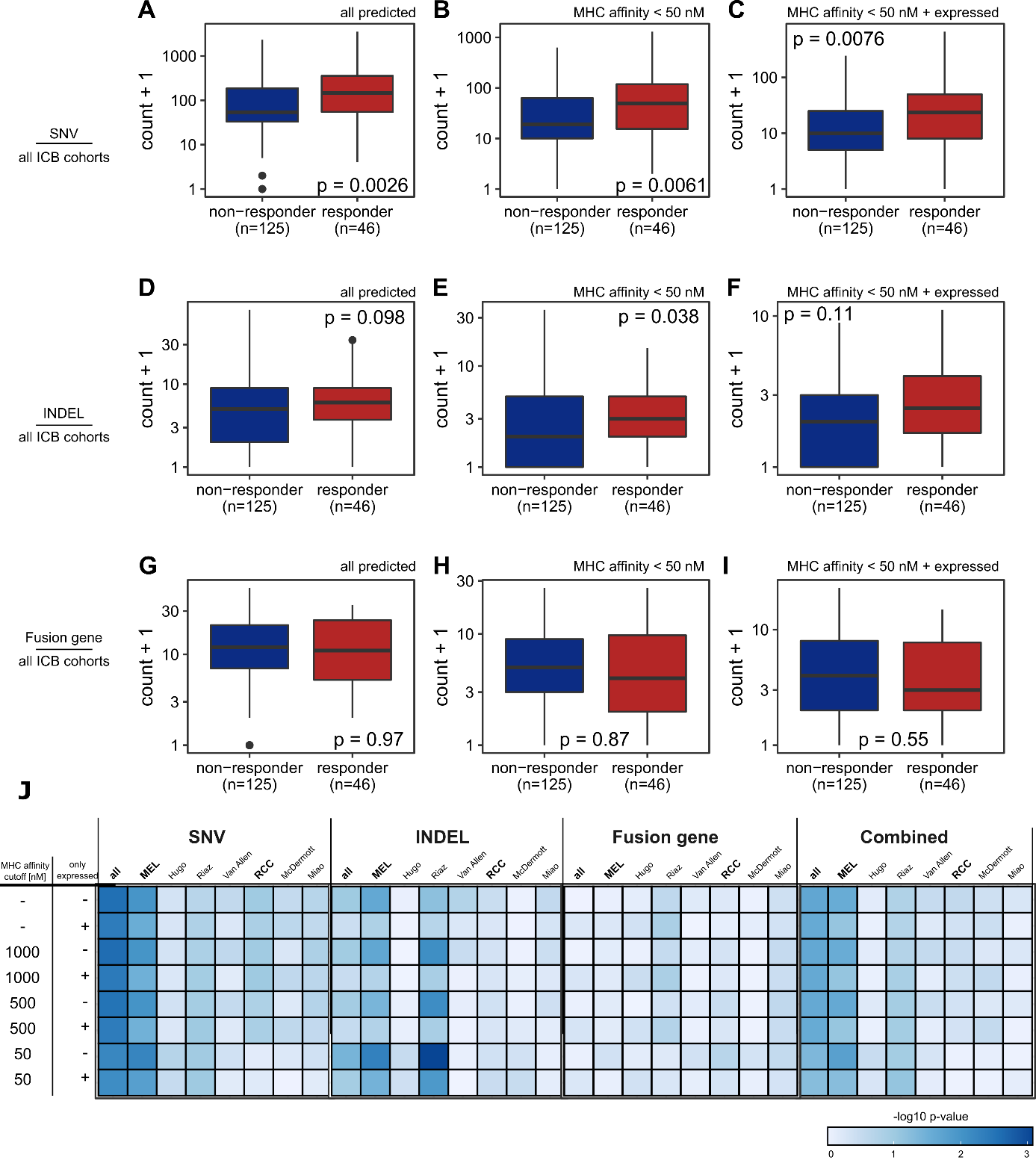
**Supplemental Figure 1: (A-C)** The SNV-derived neoantigen candidate burden was compared between responder and non-responder in a combined dataset of all ICB cohorts based on (**A**) all predicted neoantigen candidates, (**B**) candidates with MHC-I or MHC-II binding affinity < 50 nM and (**C**) expressed candidates with MHC-I or MHC-II binding affinity < 50 nM. (**D-F**) The INDEL-derived neoantigen candidate burden was compared between responder and non-responder in a combined dataset of all ICB cohorts based on (**D**) all predicted neoantigen candidates, (**E**) candidates with MHC-I or MHC-II binding affinity < 50 nM and (**F**) expressed candidates with MHC-I or MHC-II binding affinity < 50 nM. (**G-I**) The fusion gene-derived neoantigen candidate burden was compared between responder and non-responder in a combined dataset of all ICB cohorts based on (**D**) all predicted neoantigen candidates, (**E**) candidates with MHC-I or MHC-II binding affinity < 50 nM and (**F**) expressed candidates with MHC-I or MHC-II binding affinity < 50 nM. (**J**) The neoantigen candidate load was compared between responder and non-responder with respect to the mutation type, MHC binding ability and RNA expression. The heatmap represents the resulting the p-value from each comparison analogue to Figure 3D. Statistical testing was performed with Wilcoxon signed ranked test.

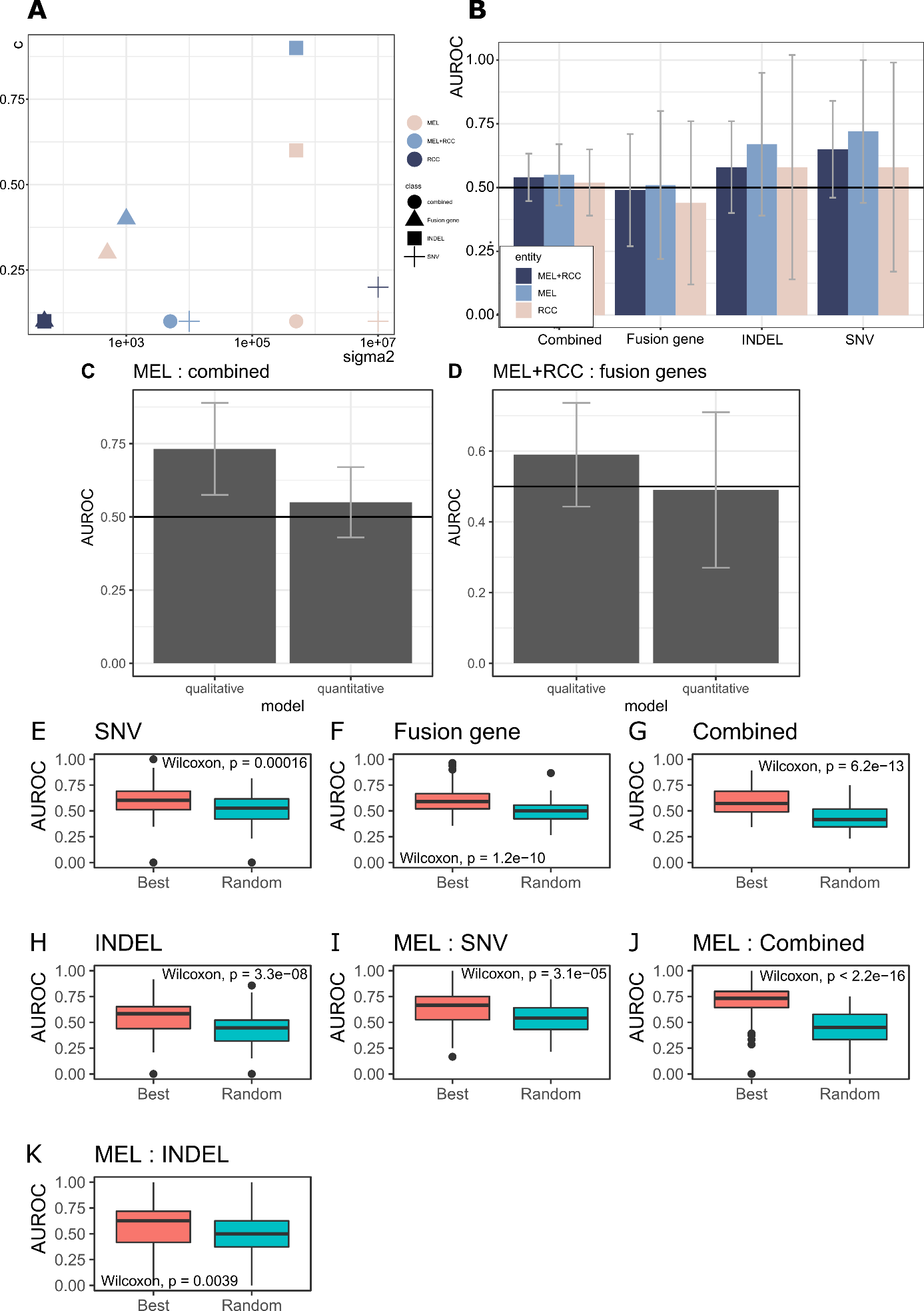

**Supplementary Figure 2**: (**A**) The optimal hyperparameter set for MILES trained on SNVs, INDELs, fusion genes or on a combined dataset of all cohorts (“MEL+RCC”), melanoma cohorts (“MEL”) and renal cell carcinoma cohorts (“RCC”). (**B**) AUROC of the neoantigen candidate load considering all predicted neoantigen candidates from SNVs, INDELs and fusion genes in a dataset of all cohorts (“MEL+RCC”), melanoma cohorts (“MEL”) and renal cell carcinoma cohorts (“RCC). Bars represent median +/- interquartile range over a nested CV. (**C-D**) Direct comparison of the qualitative and quantitative approach when (**C**) all mutation types (“combined”) are considered in the MEL cohort and if (**D**) fusion genes are considered in all cohorts (“MEL+RCC”). Bars represent median +/- interquartile range over a nested CV. (**E-K**) The optimal hyperparameter sets were used to train and evaluate the performance of MILES on a randomized dataset. Randomized refers to the randomization of neoantigen candidates across patients while keeping the original number of neoantigen candidates per patient.

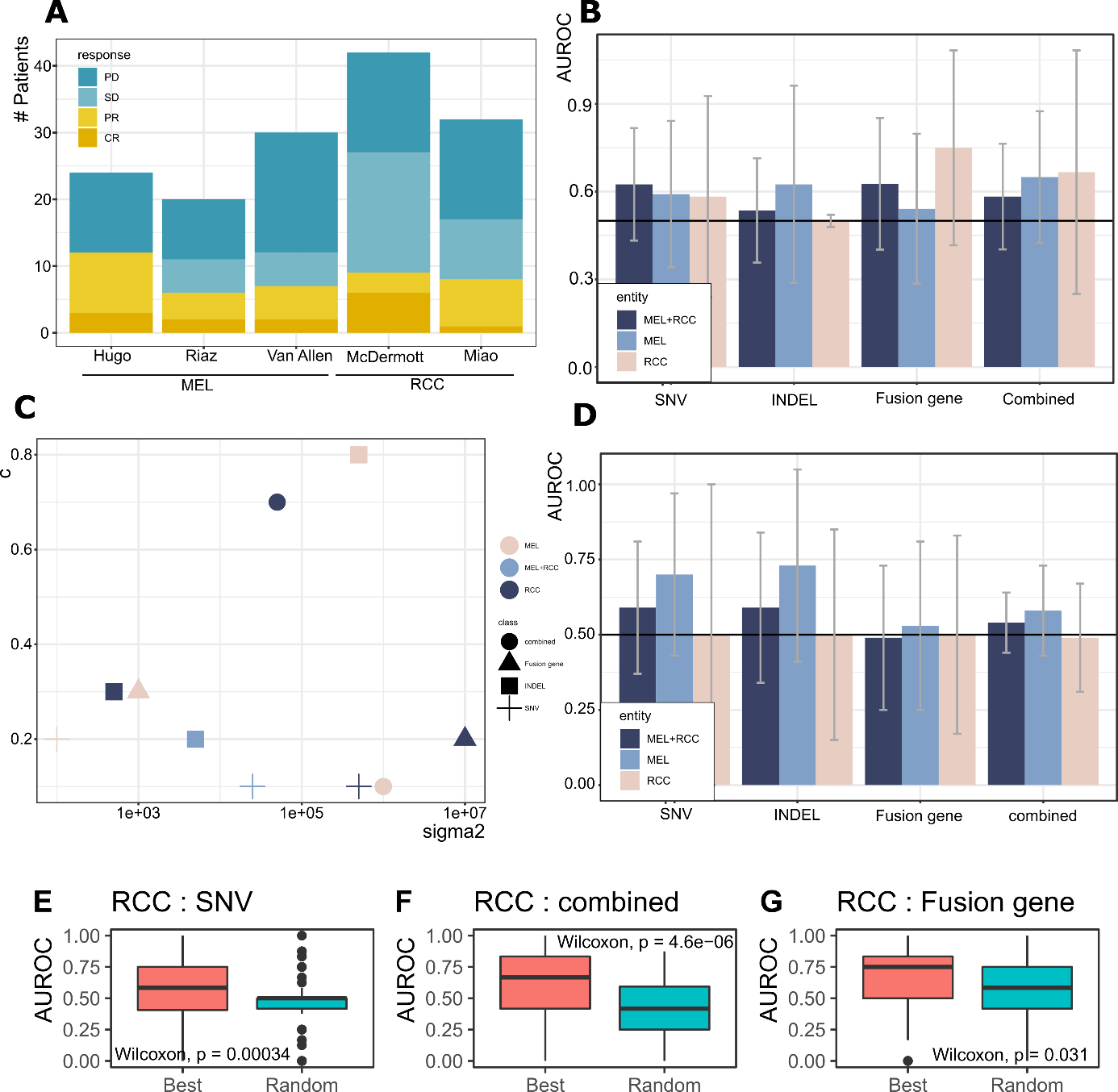

**Supplementary Figure 3**: (**A**) Distribution of response categories within the individual cohorts. (**B**) The median performance of MILES on a dataset of SNVs, fusion genes or INDELS and a dataset combining all mutation classes (“combined”) for all cohorts (“MEL+RCC”), the melanoma cohorts (“MEL”) and the renal cell carcinoma cohorts (“RCC”) if patients with stable disease (SD) are excluded from the analysis. Bars represent median +/- interquartile range over a nested CV. (**C**) The optimal hyperparameter set for MILES trained on SNVs, INDELs, fusion genes or a combined dataset in the context of the underlying cohort if patients with stable disease (SD) are excluded from the analysis. (**D**) AUROC of the neoantigen candidate load considering all predicted neoantigen candidates from SNVs, INDELs and fusion genes in a dataset of all cohorts (“MEL+RCC”), melanoma cohorts (“MEL”) and renal cell carcinoma cohorts (“RCC) without patients with stable disease. Bars represent median +/- interquartile range over a nested CV. (**E-G**) The optimal hyperparameter sets were used to train and evaluate the performance of MILEs on a randomized dataset without patients with stable disease for the RCC cohort. Randomized refers to the randomization of neoantigen candidates across patients while keeping the original number of neoantigen candidates per patient in the respective dataset.

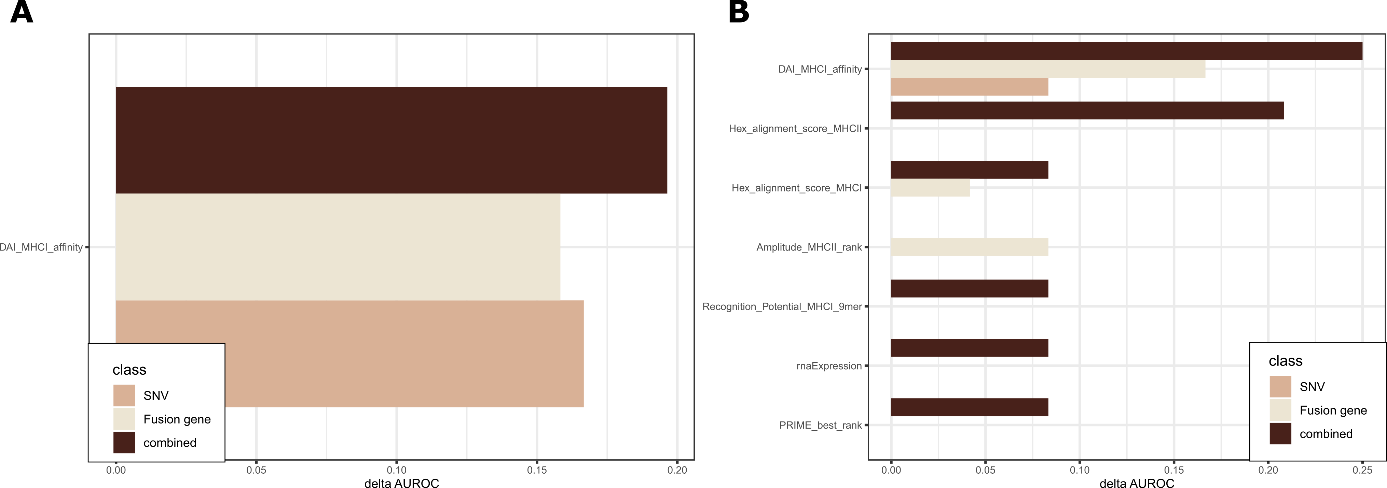

**Supplementary Figure 4**: (**A**) Feature importance for MILES on neoantigen candidates from melanoma cohorts. (**B**) Feature importance for MILES on neoantigen candidates from renal cell carcinoma cohorts, excluding patients with stable disease. Features with delta AUROC ≥0.05 in at least one approach are shown.
